# The strength of a NES motif in the nucleocapsid protein of human coronaviruses is related to genus, but not to pathogenic capacity

**DOI:** 10.1101/2020.10.06.328138

**Authors:** Maria Sendino, Miren Josu Omaetxebarria, Jose Antonio Rodriguez

## Abstract

Seven members of the *Coronaviridae* family infect humans, but only three (SARS-CoV, SARS-CoV-2 and MERS-CoV) cause severe disease with a high case fatality rate. Using *in silico* analyses (machine learning techniques and comparative genomics), several features associated to coronavirus pathogenicity have been recently proposed, including a potential increase in the strength of a predicted novel nuclear export signal (NES) motif in the nucleocapsid protein.

Here, we have used a well-established nuclear export assay to experimentally establish whether the recently proposed nucleocapsid NESs are capable of mediating nuclear export, and to evaluate if their activity correlates with coronavirus pathogenicity.

The six NES motifs tested were functional in our assay, but displayed wide differences in export activity. Importantly, these differences in NES strength were not related to strain pathogenicity. Rather, we found that the NESs of the strains belonging to the genus *Alphacoronavirus* were markedly stronger than the NESs of the strains belonging to the genus *Betacoronavirus*.

We conclude that, while some of the genomic features recently identified *in silico* could be crucial contributors to coronavirus pathogenicity, this does not appear to be the case of nucleocapsid NES activity, as it is more closely related to coronavirus genus than to pathogenic capacity.

## INTRODUCTION

Seven members of the *Coronaviridae* family (SARS-CoV, SARS-CoV-2, MERS-CoV, HCoV-NL63, HCoV-229E, HCoV-HKU1 and HCoV-OC43) are known to infect humans, but only the first three cause severe disease. Identifying molecular determinants of coronavirus pathogenicity is an important issue. In this regard, several genomic features that could differentiate highly pathogenic coronaviruses from less pathogenic strains have been recently identified *in silico*, using machine learning techniques and comparative genomics (1).

Eleven genomic regions corresponding to four different viral proteins, including the nucleocapsid (N) protein, were found to predict a high pathogenic capacity, but the underlying biological mechanisms remain to be elucidated. Interestingly, pathogenicity-associated deletions, insertions and substitutions within the N protein mapped to four potential nucleocytoplasmic transport motifs: three nuclear localization signals (NLSs) and one nuclear export signal (NES). In highly pathogenic strains, these four motifs showed an increased content of positively charged amino acids and were proposed to have an enhanced transport activity (1). However, while a direct correlation between NLS activity and positive charge has indeed been reported (2), such a correlation cannot be extrapolated to the NES motif. The activity of NES motifs, (i.e. their ability to bind to the nuclear export receptor CRM1) crucially relies on the presence of several hydrophobic (Φ) residues with a characteristic spacing (Φ_0_-X_(2)_-Φ_1_-X_(2-3)_-Φ_2_-X_(2-3)_-Φ_3_-X-Φ_4_, where X represents any amino acid) (reviewed in 3). Importantly, this NES “consensus” pattern is remarkably loose (4), and thus, predicting NES activity solely from amino acid sequence is a notoriously challenging task (5).

A different, more carboxy-terminally located sequence in the N protein of SARS-CoV was previously reported as a functional NES (6), but the more recently predicted motif (figure 1A) has not been yet experimentally evaluated. Two necessary steps towards elucidating the mechanism behind a potential relationship between the recently proposed nucleocapsid NESs and coronavirus pathogenicity are (i) to establish if these sequence motifs are functional (i.e, capable of mediating CRM1-dependent nuclear export) and, in that case, (ii) to determine if their activity level (their strength) correlates with the pathogenic capacity of the corresponding viral strain. To address these issues, we experimentally tested the NES motifs predicted in the N protein of SARS-CoV-2, MERS-CoV, HCoV-NL63, HCoV-229E, HCoV-HKU1 and HCoV-OC43 strains (Figure 1B) using a well-established nuclear export assay (7). Our results validate these sequences as functional NES motifs with different levels of nuclear export activity, and show that differences in NES strength are related to genus (*Alpha*- or *Betacoronavirus*), but not to pathogenicity of human coronavirus strains.

**Figure 1.**
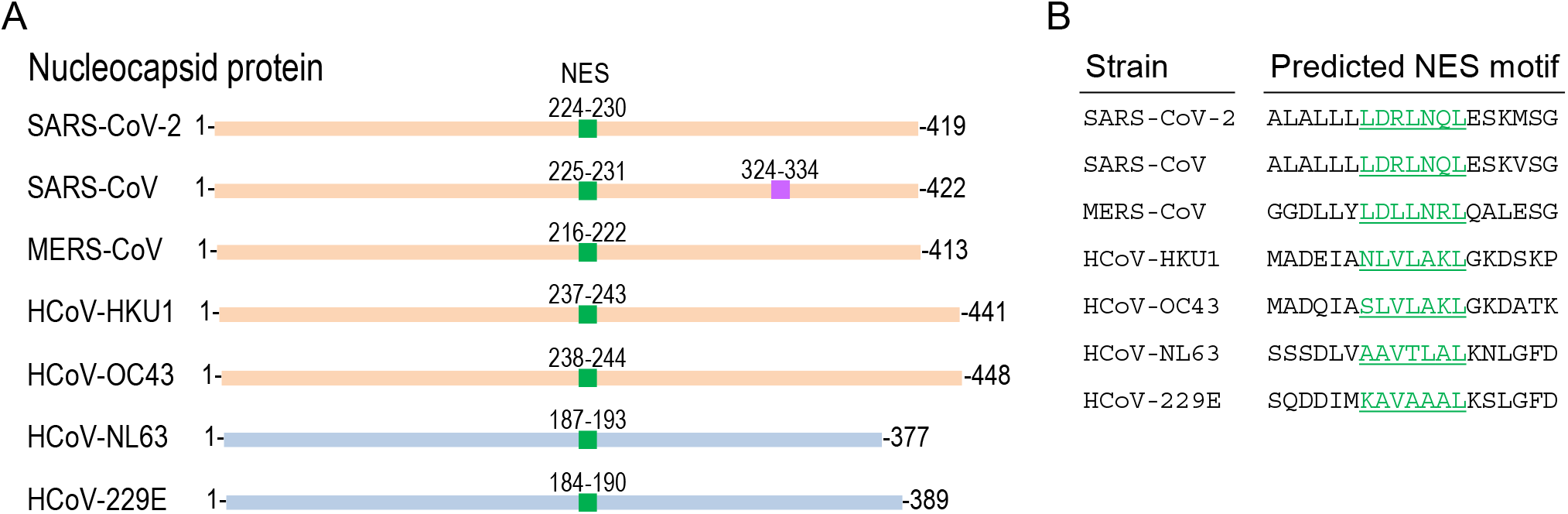
Location and amino acid sequence of predicted NES motif in the nucleocapsid (N) protein of human coronaviruses. **A**. Schematic representation (not at scale) of the nucleocapsid (N) protein of human coronaviruses indicating the location of the novel predicted NES motifs (green) whose activity has been proposed to correlate with strain pathogenicity (1). The location of a previously reported NES motif in SARS-CoV N protein (6) is shown in purple. Proteins corresponding to strains of the *Betacoronavirus* genus are represented in orange, and those corresponding to strains of the *Alphacoronavirus* genus are represented in blue. **B**. Amino acid sequence of a segment of the N protein from different coronavirus strains encompassing the proposed NES described in (1) (highlighted in green). These 19 amino acid-long sequences were tested using the Rev(1.4)-GFP nuclear export assay (7). Given the high similarity between SARS-CoV and SARS-CoV-2 motifs, only the last one was tested.

## MATERIALS AND METHODS

### Cloning procedures

The Rev(1.4)-GFP plasmid (7) was a gift from Dr. Beric Henderson (University of Sydney, Australia). DNA sequences encoding coronavirus nucleocapsid NES motifs were purchased as gBlocks (IDT), digested with BamHI and AgeI and cloned into Rev(1.4)-GFP. Constructs were confirmed using DNA sequencing (StabVida).

### Cell culture and transfection

HeLa cells were grown in Dulbecco’s modified Eagle’s medium (DMEM) supplemented with 10% fetal bovine serum (FBS), 100 U/ml penicillin and 100 μg/ml streptomycin (all from Gibco; ThermoFisher Scientific) at 37°C in a humidified atmosphere containing 5% CO_2_. 24 hours before transfection, cells were seeded in 12-well plates with glass coverslips. Transfections were carried out using X-tremeGENE 9 DNA transfection reagent (Roche Diagnostics) following manufacturer’s instructions.

### Rev(1.4)-GFP nuclear export assay

Rev(1.4)-GFP plasmids with the different NES motifs (as well as the Rev(1.4)-GFP plasmid, as negative control) were transfected into duplicated wells of HeLa cells. 24 hours after transfection, 10 μg/ml cycloheximide (CHX) was added to all the wells to arrest protein translation and thus ensure that any fluorescent signal present in the cytoplasm corresponds to exported and not to newly synthesized GFP-tagged proteins. For each transfection, the cells in one of the wells were additionally treated with 5 μg/ml actinomycin D (AD) to block nuclear import mediated by Rev NLS and thus facilitate detection of weaker NESs (7). Both drugs were purchased from Sigma-Aldrich. After 3 hours of incubation with the drugs, cells were fixed with 3.7% formaldehyde (Sigma-Aldrich) in phosphate-buffered saline (PBS) for 30 min, washed with PBS, and mounted onto microscope slides using DAPI-containing Vectashield (Vector Laboratories). Slides were analysed using a Zeiss Axioskop fluorescence microscope. To ensure unbiased assessment, the identity of the samples was masked before the analysis. At least 200 transfected cells per sample were examined to establish the proportion of cells where the reporter shows nuclear (N), nuclear and cytoplasmic (NC) or cytoplasmic (C) localization. Based on this proportion, each of the tested motifs was assigned a nuclear export activity score using the assay scoring system (7). Representative images were taken using a Zeiss Apotome2 microscope and Zen2.6 Blue edition software.

## RESULTS AND DISCUSSION

The Rev(1.4)-GFP assay (7) is a well-established method that has been used in numerous studies to experimentally establish the functionality of potential NES motifs and to compare the strength of different NESs. With this assay, functional NES motifs can be ascribed a score ranging between 1+ (weakest) and 9+ (strongest). As shown in Figure 2, the novel predicted NES motifs in the N protein of different human coronavirus strains tested here were all functional in the export assay. Of note, the N protein of SARS-CoV has been reported before to have a more carboxy-terminally located NES (6). The contribution of these sequences to the localization of human coronavirus N proteins needs to be further investigated. On the other hand, functional NES motifs have been also previously identified in the accessory proteins 3b and 9b of SARS-CoV (8-10). The presence of functional NESs in viral proteins is a common finding, and in fact, several viruses crucially rely on CRM1-mediated nuclear export of viral components (either RNA or proteins) during various stages of their life cycle (reviewed in 11). Together with previous findings (6, 8-10), our data support the possibility that nucleocytoplasmic shuttling may play an important role, still to be determined, in the life cycle of human coronaviruses.

**Figure 2.**
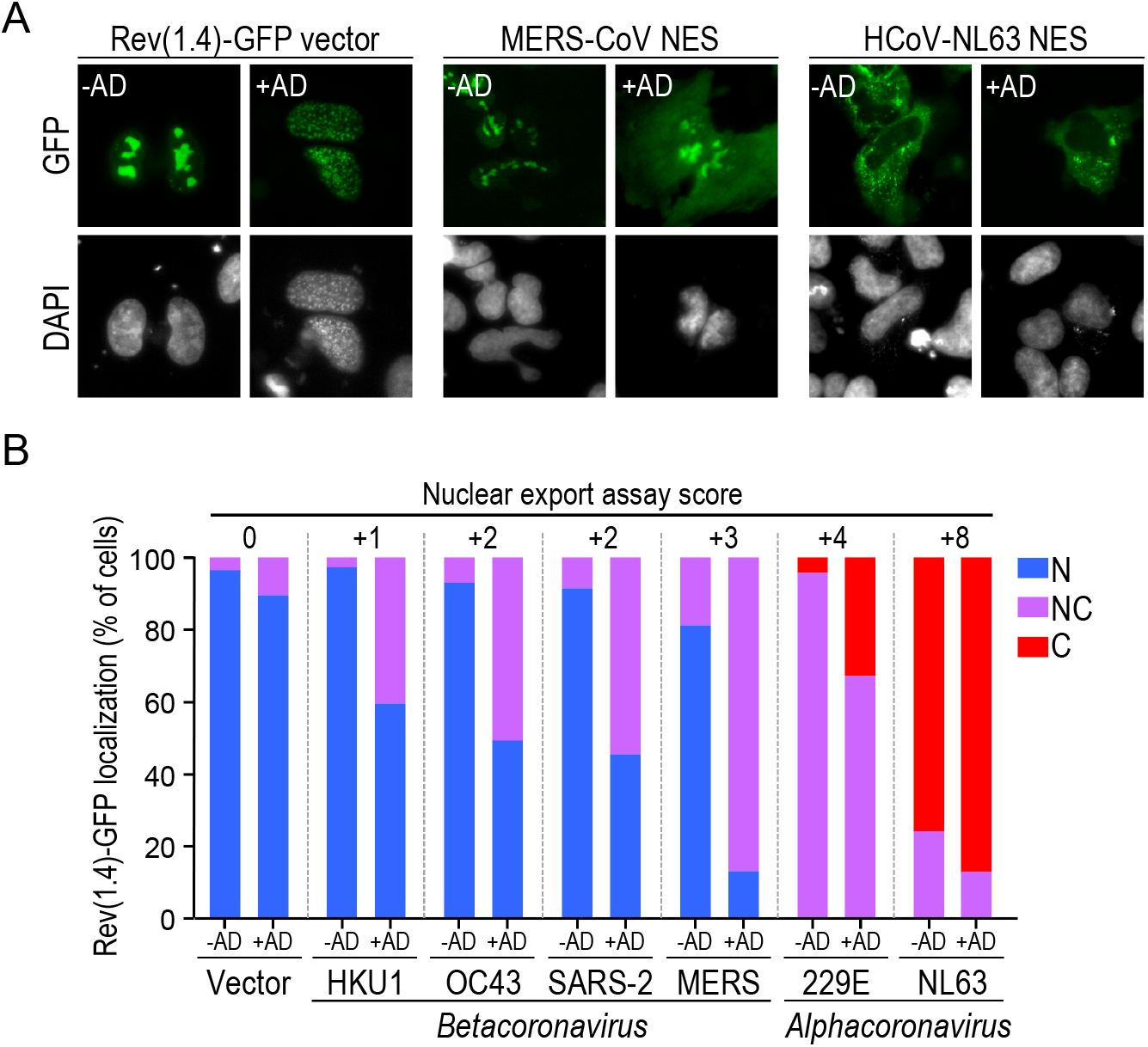
Experimental analysis of predicted NES motifs in human coronavirus N protein. **A**. Images show representative examples of the results of the nuclear export assay in HeLa cells. The empty Rev(1.4)-GFP vector was used as a negative control. Results for MERS-CoV NES and HCoV-NL63 NESs are shown. Where indicated (+AD), Actinomycin D was added to block nuclear import mediated by Rev NLS, and thus detect the activity of weaker NESs (7). Cell nuclei are visualized using DAPI. **B**. Graphs represent the percentage of HeLa cells showing nuclear (N), nuclear and cytoplasmic (NC) or cytoplasmic (C) localization of the reporter containing the indicated NES motif. At least 200 cells per sample were scored. From these percentages, each NES was assigned an export assay score as described in (7), which is indicated in the graph by the numbers above the bars.

Importantly, the six NES motifs tested here displayed a wide range of nuclear export activity (scores between 1+ and 8+). However, in contrast to what has been proposed based on *in silico* analyses (1), the differences in NES activity were not obviously related to pathogenicity (Figure 3). Rather, we found that the NESs of strains belonging to the genus *Alphacoronavirus* (HCoV-NL63 and HCoV-229E) were stronger (mean activity score= 6) than the NESs of the strains belonging to the genus *Betacoronavirus* (SARS-CoV-2, MERS-CoV, HCoV-HKU1 and HCoV-OC43; mean activity score= 2).

**Figure 3.**
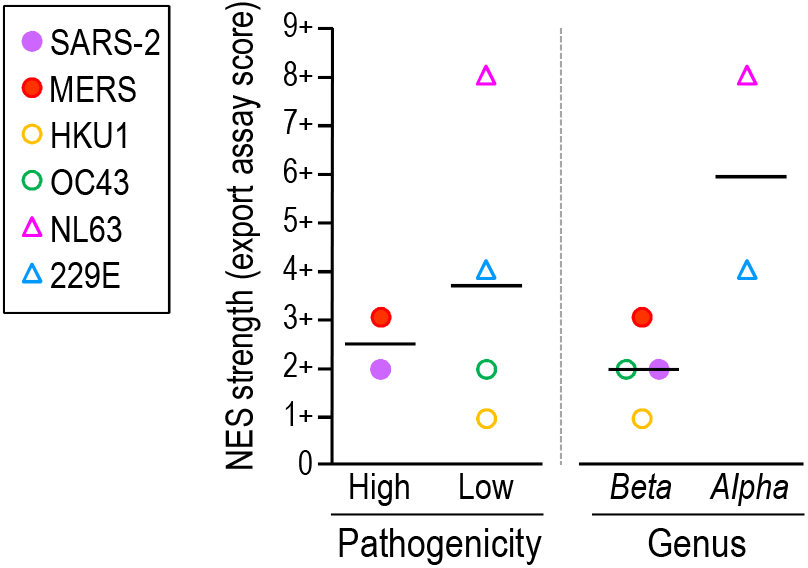
Relationship between nucleocapsid NES strength and pathogenic capability or genus of human coronaviruses. Graphs represent the strength (nuclear export assay score) of the nucleocapsid NESs in relation to the pathogenicity (left) or the genus (right) of the corresponding viral strain. Horizontal bars represent the mean activity of each group.

## CONCLUSION

We conclude that, while some of the features recently identified *in silico* (1) could be crucial contributors to coronavirus pathogenicity, this is not the case of the activity of the nucleocapsid NES motifs tested here, as it seems more closely related to coronavirus genus than to pathogenic capacity.

## ACKNOWLEDGMENTS

We are grateful to Beric Henderson for the generous gift of the Rev(1.4)-GFP plasmid. This work was supported by grants from the Spanish Government MINECO-FEDER (SAF2014-57743-R), the Basque Country Government (IT1257-19) and the University of the Basque Country (UFI11/20), as well as a fellowship from the Basque Country Government (to M.S.).

